# Behavioral and neural evidence for perceptual predictions in social interactions

**DOI:** 10.1101/2024.11.06.622031

**Authors:** Juanzhi Lu, Lars Riecke, Beatrice de Gelder

## Abstract

The ability to predict others’ behavior is crucial for social interactions. The goal of the present study was to test whether predictions are derived during observation of social interactions and whether these predictions influence how the whole-body emotional expressions of the agents are perceived. Using a novel paradigm, we induced social predictions in participants by presenting them with a short video of a social interaction in which a person approached another person and greeted him by touching the shoulder in either a neutral or an aggressive fashion. The video was followed by a still image showing a later stage in the interaction and we measured participants’ behavioral and neural responses to the still image, which was either congruent or incongruent with the emotional valence of the touching. We varied the strength of the induced predictions by parametrically reducing the saliency of emotional cues in the video.

Behaviorally, we found that reducing the emotional cues in the video led to a significant decrease in participants’ ability to correctly judge the appropriateness of the emotional reaction in the image. At the neural level, EEG recordings revealed that observing an angry reaction elicited significantly larger N170 amplitudes than observing a neutral reaction. This emotion effect was only found in the high prediction condition (where the context in the preceding video was intact and clear), not in the mid and low prediction conditions. We further found that incongruent conditions elicited larger N300 amplitudes than congruent conditions only for the neutral images. Our findings provide evidence that viewing the initial stages of social interactions triggers predictions about their outcome in early cortical processing stages.

## 1 Introduction

Primates live in complex social networks that are built and maintained by interactions between the members. The primate brain is fine-tuned to perceive nonverbal communication signals from conspecifics. In the domain of vision, social signals are predominantly provided by movements of the face and the body, whether these are displayed by single agents or in interactions. The pioneering research by Heider and Simmel (Heider & Simmel, 1944) demonstrated that humans discern intricate details about others’ interactions based on simple movement cues. In the last two decades, cognitive and affective neuroscientists have started exploring the brain basis of the competences required to engage actively in social interactions and to understand the meaning of observed social interactions (Poyo Solanas & de Gelder, 2025). The centrality of social interaction is underscored by findings showing that an individual’s expressive postures are judged differently depending on whether they are viewed as part of an interaction with another individual. Using well-controlled computer animations, Christensen et al (2024) showed that the emotional expression of an individual agent is perceived differently when the agent is shown in isolation vs. as part of a social interaction (Christensen et al., 2024). Another behavioral study found that emotions were perceived differently in a social interaction context in which two agents interacted vs. did not interact (Abramson et al., 2021). Participants were instructed to categorize the target agent’s emotions (either fear or anger), with the other agent serving as contextual cues. It was found that recognizing fear was easier when participants interacted with an angry emotion compared to a fearful emotion. This effect was observed when participants viewed body or body-face compound stimuli, but not when they viewed faces alone. These studies indicate that body gestures and movements play an important role in emotion perception during social interaction.

Research on the neural basis of affective signals from whole-body postures and movements is still a relatively underexplored field (de Gelder, 2006; de Gelder & Solanas, 2021). Functional magnetic resonance imaging (fMRI) and electroencephalography (EEG) studies have shown that the brain is fine-tuned to details of whole-body postures and movements. Furthermore, observers are not passively registering the visual input from whole-body expressions, but the brain is actively preparing for an adaptive response, such as when a defensive reaction is called for (de Gelder et al., 2004). Importantly, for many familiar actions, once the goals of the action are understood, the end stages can be successfully predicted, as shown in studies comparing basketball novices vs. experts. The latter needed less information to accurately predict where a ball was going to land (Abreu et al., 2012; Özkan et al., 2019). This ability to predict the outcome of an ongoing action is especially relevant when we observe two agents in the course of a social interaction (McMahon & Isik, 2023). One study used short video clips of real-life interactions between dyads and asked participants to predict the outcome of the observed social interaction (Epperlein et al., 2022). They found that participants predicted the outcome of a social interaction less accurately in an aggressive context compared to a playful or neutral context, indicating that predictions depend on the emotional information available during observations of social interactions.

A few studies have examined how prediction operates in the course of neural processing of emotional stimuli (Baker et al., 2023; Vogel et al., 2015). For example, Baker et al. (2023) found that N170 and N300 responses to face stimuli are sensitive to emotion-prediction errors, showing stronger responses to unpredictable facial emotional expressions than predictable ones. Similarly, Vogel et al. (2015) found that the mismatch negativity (MMN), a mid-latency event-related potential (ERP) component thought to reflect regularity violations, is sensitive to prediction errors based on facial emotional expressions. Their study showed that incongruent emotional faces (e.g., a neutral face followed by a fearful face) elicited larger MMN amplitudes compared to congruent faces (e.g., a neutral face followed by another neutral face). Another ERP study found that perceiving two consecutive emotional expressions elicits a stronger N400 response when the two expressions are incongruent rather than congruent (Calbi et al., 2017). This effect was observed regardless of whether the expression was conveyed by still images of the face or the body, and it might hint at a prediction error response.

Other studies have focused on the N170, as it is linked to the processing of not only faces but also bodies (Baker et al., 2023; Calbi et al., 2017; He et al., 2018; Stekelenburg & de Gelder, 2004; Van Heijnsbergen et al., 2007). Some studies have found effects of emotional expression on the body-evoked N170 (Lu et al., 2023), while others have not (Stekelenburg & de Gelder, 2004; Van Heijnsbergen et al., 2007). Given the previous observation of an emotion-prediction effect on the face-evoked N170 (Baker et al., 2023), it is still an open question whether the body-evoked N170 is affected by emotion predictions when observing social interactions. Taken together, the N300 and N400 may serve as neural markers of violations of higher-order visual predictions, whereas the N170 may specifically reflect the visual processing of bodies.

We hypothesized that: 1) Observers of a social interaction derive predictions from their observations about the outcome of the interaction; and 2) These putative social predictions automatically and rapidly influence how the outcome of the ongoing social interaction is perceived. We tested our hypotheses with a novel paradigm: Participants watched a short video clip of a social interaction between two agents, in which agent A approached agent B and touched him on the shoulder, whereupon agent B turned around to face agent A. The videos were stopped before the end and then followed a by a still probe image, which was the final frame of the full clip disclosing agent B’s reaction to the interaction. In the perceptual task, participants judged the appropriateness of the agents’ reaction from the agent’s bodily expression. For the neural measures, we focused on the ERP components N170, N300, and N400, as reviewed above. By presenting the video clip prior to the still probe we could temporally separate the putative prediction effects of the video from its (shorter-lived) sensory effects. To investigate the impact of social prediction on observing social interactions, we varied both the strength and the correctness of the predictions that observers could derive from the clip. Prediction strength was varied across three levels as follows: in the main “high prediction” condition, the video clearly showed how agent A approached and touched agent B. In the “mid prediction” condition, social interaction information was reduced by backward presentation of the video. Finally, in the “low prediction” condition, each video frame was scrambled, effectively removing any social cues from the video and preventing emotion prediction.

Prediction correctness, referred to below as prediction error, was varied by manipulating the emotional congruence between the probe image and the preceding video. This was implemented by preceding each probe condition (image of a neutral or angry reaction; see above) with either a “neutral” video (in which agent A gently touched agent B’s shoulder) or an “angry” video (in which agent A abruptly pulled agent B’s shoulder). The incongruent condition was designed to trigger prediction errors in participants.

We expected that: 1) If observers of a social interaction derive predictions from it about its outcome, our participants should show more accurate responses in the perceptual task when the preceding clip allows for stronger predictions. 2) If these social predictions influence the processing of the ongoing social interaction, our participants should show neural changes in response to the probe. Specifically, body-related responses (N170) and prediction-related responses (N300 and N400) should reflect variations in prediction strength and prediction errors.

## 2 Methods

### 2.1 Participants

Thirty healthy participants were recruited from the student population at Maastricht University. Two participants’ data were rejected because one participant did not follow the task instructions and another participant’s ERPs data (N170, N300 and N400) exceeded 3 standard deviations (SD) above the mean. Twenty-eight participants’ data were included in the analysis (aged 19-34 years, 24.0 **±** 4.9 (mean ± SD); 15 male and 14 female; one left-handed). All participants had normal or corrected-to-normal vision, and no history of brain injury, psychiatric disorders, or current use of psychotropic medication. Before the experiment, participants provided written consent. They received compensation of 7.5 Euros or one study credit point for their participation. The Ethics Committee of Maastricht University approved the study, and all procedures adhered to the principles outlined in the Declaration of Helsinki (approval number: OZL_263_16_02_2023).

### 2.2 Stimuli

The stimuli consisted of video clips of social interactions and still images extracted from the end section of the videos. The videos showed a person on the right (agent A) approaching a person on the left (agent B). At the onset, agent B had his/her back turned away from agent A. Agent A approached and touched agent B on the shoulder whereupon agent B reacted to this by turning around toward agent A.

The video recordings were made with ten actors (six females and four males) who were combined to create five gender-matched pairs. For each actor pair, five “angry” social interactions and five “neutral” social interactions were recorded, resulting in ten videos per pair (50 videos in total). The still images were created by taking the last frame of the video. These images served as the probes for the participants’ task, which was to rate whether the reaction of agent B (to the touch by agent A) was appropriate. The images and videos were processed using Adobe Premiere Pro and all faces were blurred. Videos and still images were presented on a black background (size: 1150×1088 pixels), covering approximately 15×13 degrees of the participants’ visual angle in the experiment. To ensure that participants focused on the interaction between the two actors, they were instructed to fixate a white fixation cross placed at the center of the screen, located between the two actors.

### 2.3 Experimental design and procedure

Each trial started with a 1000-ms fixation period, followed by the presentation of the video. After a short gap (400-500ms) during which the screen was blank, the probe image was presented for 1000ms revealing agent B’s reaction. Subsequently, participants were instructed to answer the following question, which was shown on the screen: “Does the reaction of the person on the left match the action of the person on the right?”. Participants chose one of two response alternatives (“I guess yes” and “I guess no”) during this response interval, which lasted 2000ms (Fig 1A).

**Fig. 1.**
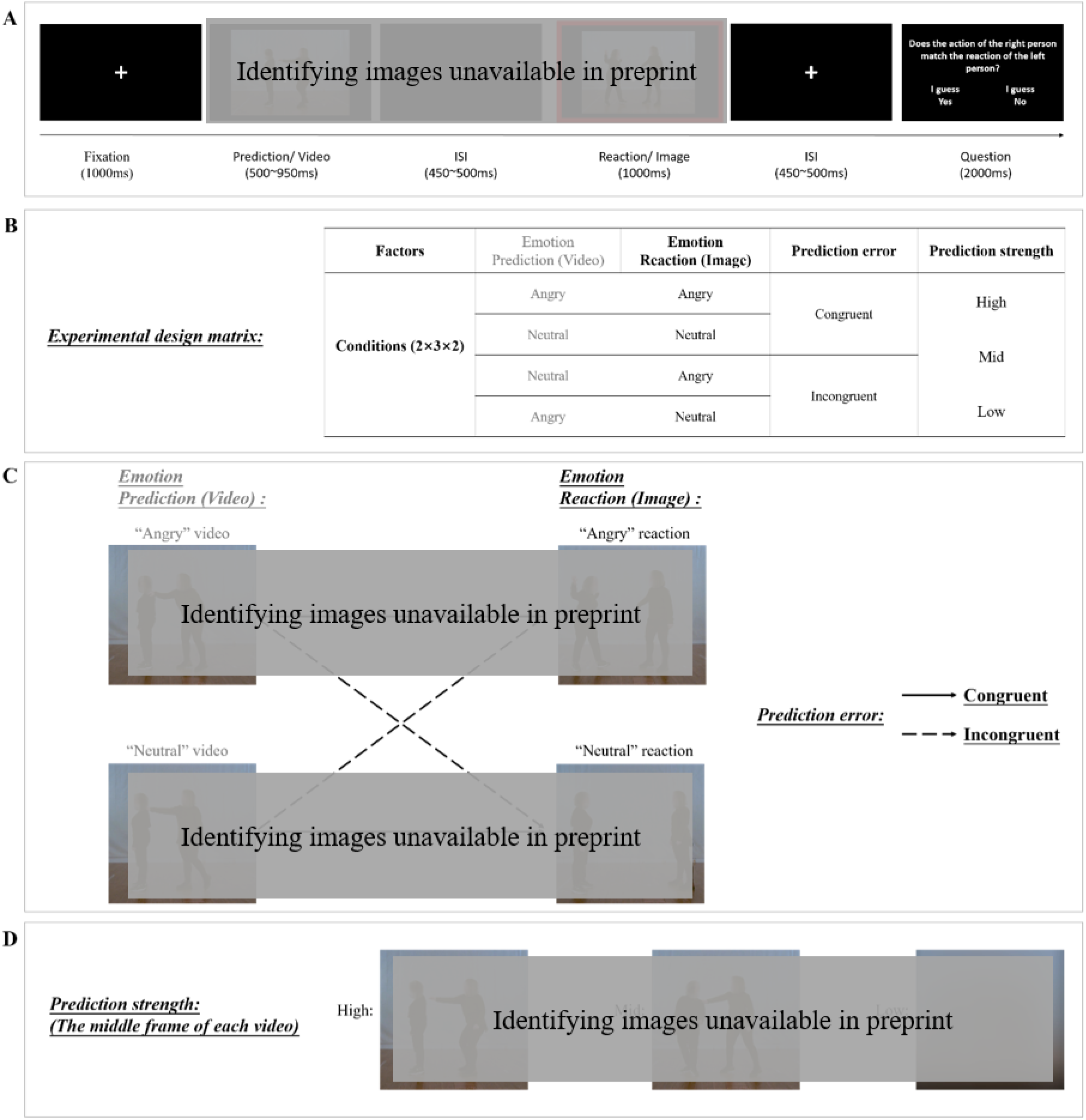
A-D. Experimental design. (A) Trial procedure. Participants watched a social interaction video followed by a still probe image. At the end of each trial, participants responded to the question on the screen by pressing one of two buttons (yes/no). ERP analysis was time-locked to the still image, see red rectangle. (B) Experimental design matrix. The study used a 2 × 3 × 2 within-subject design with factors *emotion reaction* (angry, neutral), *prediction strength* (high, mid, low), and *prediction error* (congruent, incongruent). (C) Examples of *emotional reaction* that are included in the matrix of *prediction error*. The left column of figures shows the middle frame of the “angry” video and the “neutral” video. The right column of figures shows the *emotional reaction*: the “angry” reaction and the “neutral” reaction. The solid arrows indicate congruent conditions: an angry reaction preceded by an angry video or a neutral reaction preceded by a neutral video. The dashed arrows indicate incongruent conditions: an angry reaction preceded by a neutral video or a neutral reaction preceded by an angry video. (D) Examples of prediction strength in the video, showing the first frame of high, mid and low conditions.

An example of the probe image in the two *emotion reaction* conditions (angry reaction or neutral reaction) is shown in Figure 1C. The different *prediction strength* conditions are illustrated in Figure 1D. This manipulation was implemented by playing the video either normally (high prediction), or as time-reversed or scrambled versions. In the backwards videos (mid prediction), the visibility of the actors’ movements was preserved, while the interpretation of the social action was hampered. In other words, the clip began with agent A already touching agent B’s shoulder, then releasing the hand, and finally walking away backwards (from left to right). In the scrambled videos (low prediction), each frame was masked with Gaussian masks so that both movement and social action information were largely reduced (Figure 1D).

The manipulation of *prediction error* was implemented by pairing each video clip with either its original last frame (*congruent* condition: angry video followed by angry image, or neutral video followed by neutral image) or the last frame of the clip in which the same actors exhibited the other emotion (*incongruent* condition: angry video followed by neutral image, or neutral video followed by angry image). Example frames from the neutral and angry videos are shown in Figure 1C. Participants’ “Yes” responses on congruent trials and “No” responses on incongruent trials were considered as correct, whereas “No” responses on congruent trials and “Yes” responses on incongruent trials were considered as incorrect.

The study used a fully balanced 2 × 3 × 2 within-subject design. As described above, the first factor was *emotion reaction* (angry or neutral), the second factor was *prediction strength* (high, mid, or low), and the third factor was *prediction error* (emotional valence of image and video: congruent or incongruent). Each of the twelve conditions was presented in 25 unique trials, resulting in a total of 300 trials that were randomly presented in 4 runs, each lasting 7 minutes. Participants took a short break after the first two runs. Before the experiment, participants practiced the task on 24 trials. The whole experiment lasted around 28-35 minutes.

### 2.4 EEG acquisition

EEG data were recorded using an elastic cap with 64 electrodes placed according to the international 10-20 system and sampled at a rate of 1000Hz (BrainVison Products, Munich, Germany). Electrode Cz was used as the reference during recording and the forehead electrode (Fp1) was used as a ground electrode. Four electrodes were used to measure the electrooculogram (EOG). Two of them were used as vertical electrooculograms (VEOG). One was placed above the right eye, and another was placed below the right eye. The other two electrodes were used as a horizontal electrooculogram (HEOG), with one placed at the outer canthus of the left eye, and the other at the outer canthus of the right eye. The remaining 60 electrodes included FPz, AFz, Fz, FCz, CPz, Pz, POz, Oz, AF7, AF8, AF3, AF4, F7, F8, F5, F6, F3, F4, F1, F2, FC5, FC6, FC3, FC4, FC1, FC2, T7, T8, C5, C6, C3, C4, C1, C2, TP9, TP10, TP7, TP8, TP9, TP10, CP5, CP6, CP3, CP4, CP1, CP2, P7, P8, P5, P6, P3, P4, P1, P2, PO7, PO8, PO3, PO4, O1, and O2. Impedances for reference and ground were maintained below 5kOhm and for all other electrodes below 10kOhm.

### 2.5 EEG data preprocessing

EEG data were preprocessed and analyzed using FieldTrip version 20220104 (Oostenveld et al., 2011) in Matlab R2021b (MathWorks, U.S.). Recordings were first segmented into epochs from 500ms pre-stimulus (i.e., before the onset of the probe image) to 1500ms post-stimulus and then filtered with a 0.3-30 Hz band-pass filter. EEG data at each electrode were re-referenced to the average of all electrodes. Artifact rejection was done using independent component analysis (logistic infomax ICA algorithm (Bell & Sejnowski, 1995); on average, 2.97 ± 1.08 (mean ± SD) components were visually identified as artifacts and removed per participant. Moreover, single epochs during which the EEG peak amplitude exceeded 3 SD above/below the mean amplitude were rejected. On average, 71.04% ± 9.14% of trials were preserved and statistically analyzed per participant.

### 2.6 Event-related potential analyses

The EEG analysis focused on neural responses to the probe (reaction) image. Baseline correction was applied and involved subtracting the average amplitude in the baseline interval (−200 to 0ms) from the overall epoch. Trials were averaged for each experimental condition, resulting in ERPs used for further statistical analyses, which were performed using IBM SPSS Statistics 27 (IBM Corp., Armonk, NY, USA). We spatially separated the EEG electrodes into a temporal cluster (P7, P8, TP7, TP8, TP9, TP10) and central cluster (FCz, FC1, FC2, Cz, C1, C2, CPz, CP1, CP2), and averaged the channels within each cluster. For each cluster, we pooled all conditions and visually identified a prominent ERP component based on visual inspection of the overall ERP waveform, topographical distribution of grand-averaged ERP, and previous studies (Chen et al., 2022; Hietanen et al., 2014). The identified ERP components and their associated time windows were as follows: N170 (180-230ms) in the temporal cluster, N300 (250-350ms) in the central cluster, and N400 (350-500ms) in the central cluster. The mean amplitude was computed as the average of all electrodes within each cluster for the specific time window.

A repeated-measures 2× 3× 2 ANOVA (*Emotion reaction*: angry/neutral; *Prediction strength*: high/mid/low; *Prediction error*: congruent/incongruent) was applied to the mean amplitudes; this was done for each ERP component separately. Degrees of freedom for F-ratios were corrected with the Greenhouse–Geisser method. Bonferroni’s method was used to correct for multiple comparisons. Statistical results were considered as significant given a p-value < 0.05.

## 3 Results

### 3.1 Behavior

To verify whether our manipulation of the video induced variations in participants’ predictions, we examined the effects of prediction strength (high, mid and low) and prediction error (congruent and incongruent) on response accuracy (proportion of correct responses), pooled across *emotion reaction*. We found that the main effect of prediction strength was significant (*F* (2, 54) = 49.23, *p* < 0.001, *η_p_^2^* = 0.65) (high vs. mid: *t* (27) = 6.91, *p* < 0.001; high vs. low: *t* (27) = 8.89, *p* < 0.001; mid vs. low: *t* (27) = 4.85, *p* < 0.001). Accuracy was highest for the high prediction condition (0.77 ± 0.15), followed by the mid prediction condition (0.68 ± 0.15), and lowest for the low prediction condition (0.54 ± 0.06). These findings indicate that our manipulation of contextual information was effective: reducing the amount of information in the preceding video led to a decrease in prediction accuracy. We found that the main effect of prediction error was not significant (*F* (1, 27) = 0.39, *p* = 0.536, *η_p_^2^* = 0.01), suggesting that task difficulty did not differ significantly between congruent (0.68 ± 0.14) and incongruent (0.65 ± 0.16) conditions.

To test whether participants’ choices/accuracy were above chance level, we conducted a one-sample t-test comparing participants’ accuracy in each prediction strength (high/mid/low) and prediction error (congruent/incongruent) condition vs. 0.5. The accuracy in all conditions was significantly above chance level (*ps* < 0.002).

**Fig. 2.**
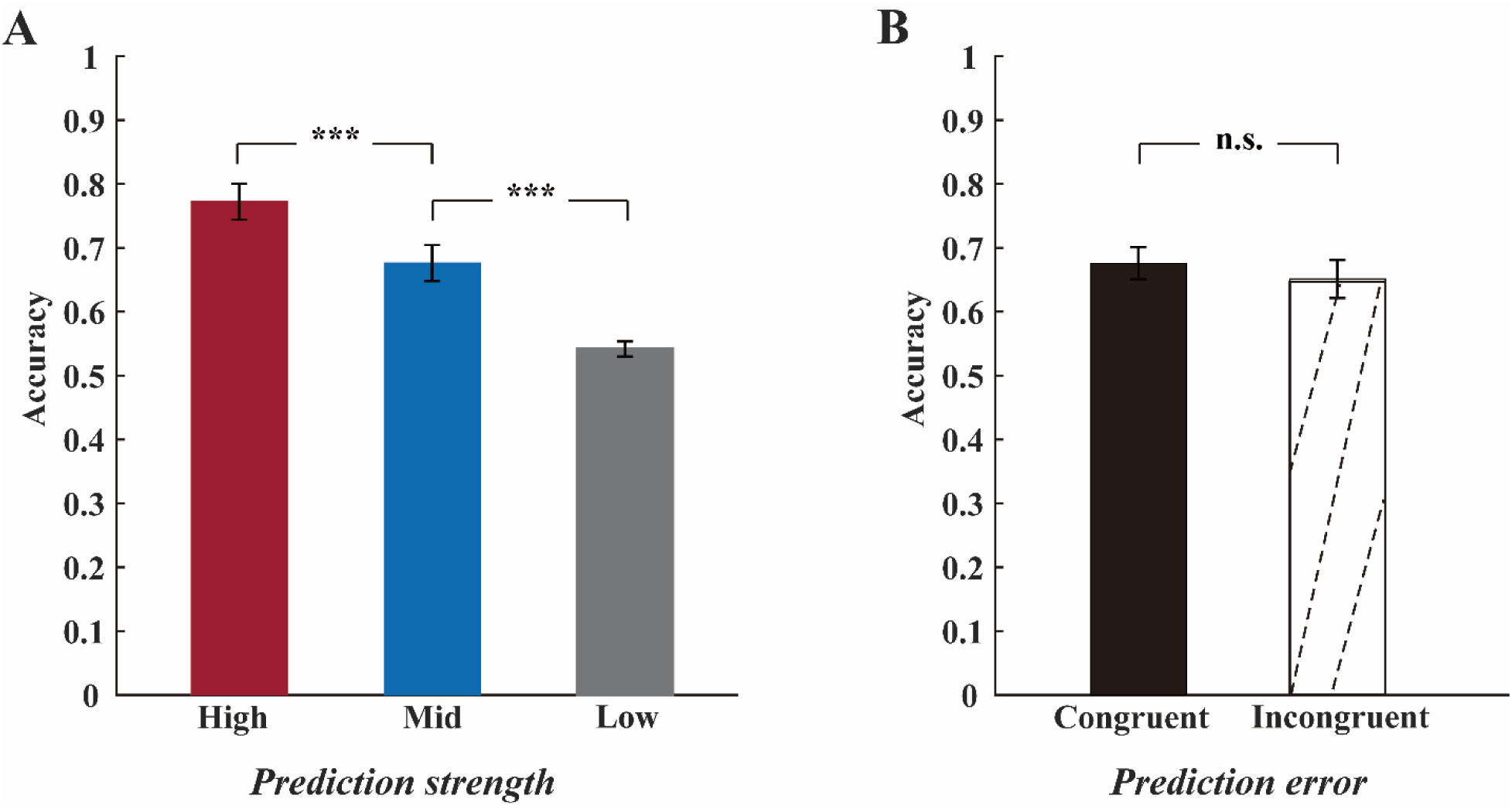
(A) Means and standard error (SE across participants) of accuracy per prediction strength condition (high, mid and low). (B) Means and SE of accuracy per prediction error condition (congruent and incongruent). ***: *p* <0.001, n.s.: non-significant.

### 3.2 ERPs

Our hypothesis concerned the effect of emotional valence (*emotion reaction*) and its modulation by contextual factors (*prediction strength* and *prediction error*). Before testing for a main effect of *emotion reaction* and its interaction with *prediction strength* and *prediction error*, we assessed the three-way interaction (*emotion reaction × prediction strength × prediction error*). This was found to be non-significant for each ERP component (N170: *F* (2, 54) = 2.44, *p* = 0.097, *η_p_^2^* = 0.08; N300: *F* (2, 54) = 0.56, *p* = 0.573, *η_p_^2^* = 0.02; N400: *F* (2, 54) = 0.83, *p* = 0.443, *η_p_^2^* = 0.03). Next we analyzed the two-way interactions, which revealed a significant *emotion reaction × prediction strength* interaction for N170 (*F* (2, 54) = 3.48, *p* = 0.040, *η_p_^2^* = 0.11), but not the other ERP components (N300: *F* (2, 54) = 0.92, *p* = 0.40, *η_p_^2^* = 0.03; N400: *F* (2, 54) = 0.18, *p* = 0.83, *η_p_^2^* = 0.01), and a significant *emotion reaction × prediction error* interaction for N300 (*F* (1, 27) = 6.47, *p* = 0.017, *η_p_^2^* = 0.19), but not the other ERP components (N170: (*F* (1, 27) = 0.05, *p* = 0.829, *η_p_^2^* = 0.00), N400: (*F* (1, 27) = 0.11, *p* = 0.745, *η_p_^2^* = 0.00)). These results are in line with our hypothesis. However, unlike hypothesized, we found no significant *prediction strength × prediction error* interaction for any ERP component (N170: *F* (2, 54) = 2.05, *p* = 0.146, *η_p_^2^* = 0.07; N300: *F* (2, 54) = 1.32, *p* = 0.28, *η_p_^2^* = 0.05; N400: *F* (2, 54) = 0.83, *p* = 0.44, *η_p_^2^* = 0.03). In the following sections, we investigated the nature of the observed interactions by testing for simple effects of the interacting factors. We also explored main effects of the factors that showed no significant interactions; these effects were not a focus of the current study and therefore the results are presented in the supplementary data.

#### Interaction effect of emotion reaction × prediction strength on N170

To disentangle the observed interaction effect of *emotion reaction × prediction strength* on N170, we analyzed simple effects of *emotion reaction* (i.e., per *prediction strength*), which revealed a significant simple effect of *emotion reaction* for the high prediction condition (*t* (27) = –5.18, *p* < 0.001) as expected, but not for the mid or low prediction conditions (mid: *t* (27) = –1.41, *p* = 0.507; low: *t* (27) = –1.74, *p* = 0.277). More specifically, angry reactions (−1.45 ± 2.00 µV) elicited larger N170 amplitudes than neutral reactions (−0.52 ± 1.87 µV) in line with previous results (Lu et al., 2023), and interestingly, this enhancing effect occurred only when the images were preceded by a fully intact video (high prediction condition).

We further observed a significant simple effect of *prediction strength* for the angry reaction. Both high and mid prediction were followed by larger N170 amplitudes than low prediction when the following reaction in the probe image was angry; the difference between high and mid prediction was not significant (Angry reaction: high vs. low: t (27) = –4.51, *p* < 0.001; mid vs. lows: t (27) = –2.62, *p* = 0.014; high vs. mid: t (27) = –2.20, *p* = 0.109). Interestingly, this simple effect of prediction strength was found only for the angry reaction, not for the neutral reaction (Neutral reaction: high vs. low: t (27) = –1.76, *p* = 0.272; mid vs. low: t (27) = –1.88, *p* = 0.214; high vs. mid: t (27) = 0.59, *p* = 1.000).

**Fig. 3.**
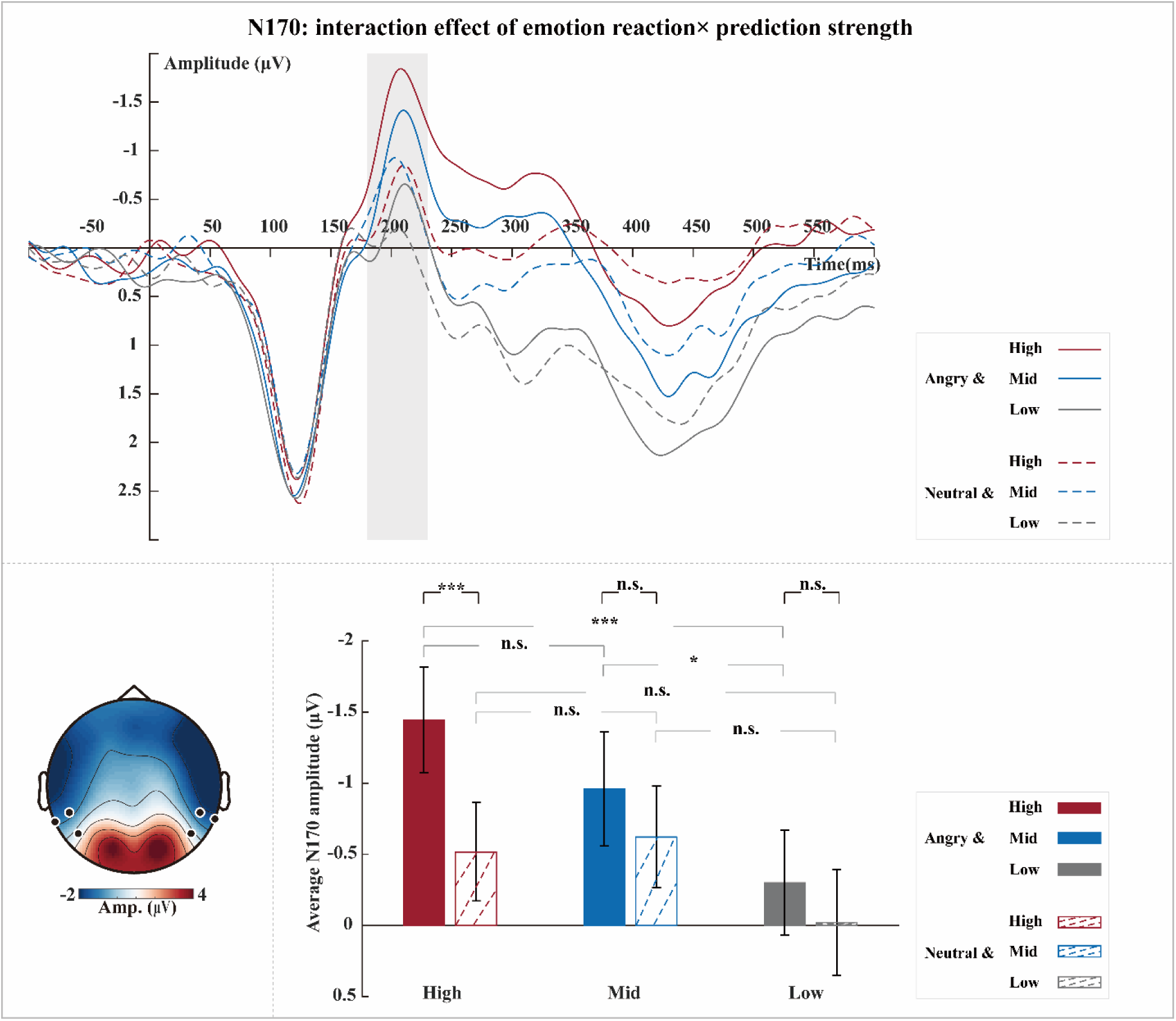
Interaction effect of *emotion reaction × prediction strength* on N170. Grand-averaged ERP waveforms of N170 per condition (angry-high, neutral-high, angry-mid, neutral-mid, angry-low, and neutral-low) (top). Waveforms were calculated by averaging the data at the electrodes P7, P8, TP7, TP8, TP9, and TP10 (see black dots in scalp map). The shaded rectangle visualizes the time window from which the average ERP amplitude was extracted (180-230ms). The topographic map was calculated by averaging the data of all conditions within a time window of 180-230ms after the onset of the probe image (bottom left). Bar plots (bottom right) illustrate the mean and SE across participants of the average N170 amplitude per condition. ***: *p* <0.001, *: *p* <0.05, n.s.: non-significant.

#### Interaction effect of emotion reaction and prediction error on N300

Further investigation of the observed interaction effect of *emotion reaction × prediction error* on N300 revealed a significant simple effect of *prediction error* for the neutral reaction (*t* (27) = 3.87, *p* = 0.001) as expected, but somewhat surprisingly not for the angry reaction (*t* (27) = –0.08, *p* = 1.000). More specifically, compared with neutral videos (−0.60 ± 1.38 µV), angry videos (-1.04 ± 1.44 µV) resulted in the subsequent neutral reaction eliciting larger N300 amplitudes.

**Fig. 4.**
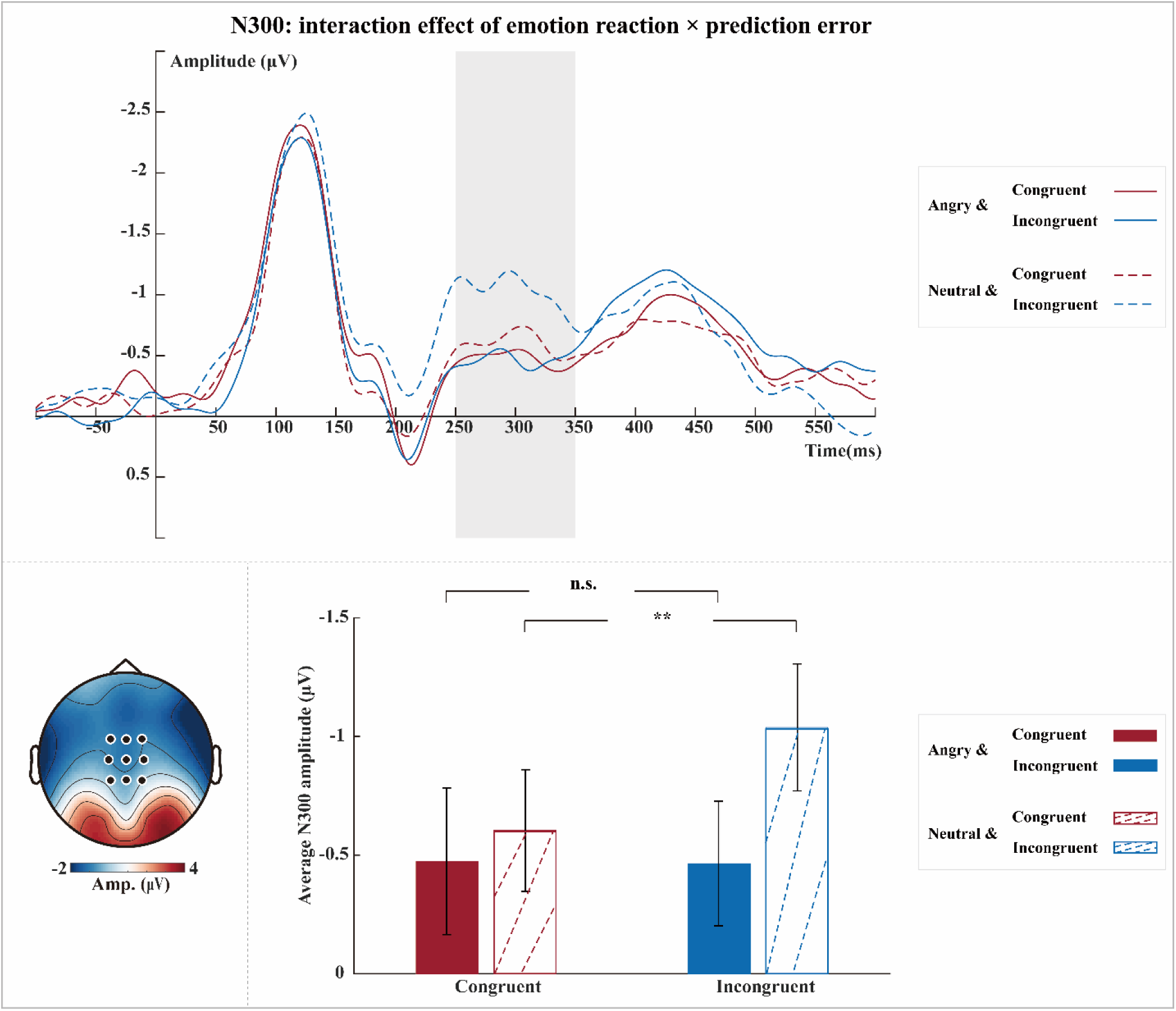
Interaction effect of *emotion reaction × prediction error* on N300. Grand-averaged ERP waveforms of N300 per condition (angry-congruent, neutral-congruent, angry-incongruent, and neutral-incongruent) (top). Waveforms were calculated by averaging the data at electrodes FCz, FC1, FC2, Cz, C1, C2, CPz, CP1, and CP2 (see black dots in scalp map). The shaded rectangle visualizes the time window from which the average ERP amplitude was extracted (250-350ms). The topographic map was calculated by averaging the data of all conditions within a time window of 250-350ms after the onset of the probe image (bottom left). Bar plots (bottom right) illustrate the mean and standard SE across participants of the average N300 amplitude per condition. ***: *p* <0.001, **: *p* <0.01, n.s.: non-significant.

## 4 Discussion

The goals of the present study were to test first, whether observers of a social interaction derive predictions about its outcome and second, whether these predictions influence how information about the outcome is processed? Our study used a novel paradigm that measures the impact of viewing the initial stages of a social interaction on how the final stages are processed. This involved manipulation of the prediction context in two different ways, by manipulating prediction strength and prediction error.

At the behavioral level, the accuracy of appropriateness judgments was highest in the high prediction condition, followed by the mid prediction condition, and lowest in the low prediction condition. Thus, our behavioral results show that participants were able to successfully judge the appropriateness of the emotional reaction (the still probe image) when the preceding video provided clear social cues (high prediction condition). Performance gradually diminished to guessing behavior when the context provided fewer emotional cues (mid and low prediction conditions). These results confirm our hypothesis that observing social interactions may lead to predictions about the outcome. At the neural level, observing an angry reaction elicited significantly larger N170 amplitudes than observing a neutral reaction. This emotion effect was only found in the high prediction condition (where the context in the preceding video was intact and clear), not in the mid and low prediction conditions. Moreover, we found that the high prediction condition elicited larger N170 amplitudes than the mid and low prediction conditions.

This prediction effect was found only in response to angry reactions. Additionally, observing social interactions can trigger prediction error effects. We found that incongruent conditions elicited larger N300 amplitudes than congruent conditions. This prediction error effect was found only in neutral reactions, not in angry reactions. Our results confirm our hypothesis that social predictions may influence the perceptual and neural processing of social interactions.

### Emotion effect on the early component N170 depends on prediction strength

Our first neural finding was that observing social interactions containing dyadic bodies evoked a clear N170 response. Previous studies have shown that the N170 is a marker of visual body processing (Borhani et al., 2015; de Gelder et al., 2004; Lu et al., 2023; Meeren et al., 2005; Stekelenburg & de Gelder, 2004). Here, we extend these previous findings by showing that the N170 is sensitive not only to a single body but also to body expressions in interactions involving two agents. Hence, our results are consistent with findings about the primacy of social interactions (Abassi & Papeo, 2020). Concerning the sensitivity of the N170, we further observed that the N170 is stronger for angry compared to neutral expressions. This is consistent with our recent finding (Lu et al., 2023) and, more importantly, extends previously observed emotional expression effect from single images and single-body expressions to social interaction situations.

Our main finding here is that the emotional expression effects during observation of interactions are only seen in the high prediction condition. In other words, neural discrimination between angry and neutral interaction images, as reflected by the N170, was not evident when the preceding social context videos did not allow emotion predictions (mid and low prediction conditions). Moreover, we found that predictions were impacted by emotional context, such that high predictability elicited larger N170 amplitudes than lower predictions for videos of angry body interactions. This result indicates that the N170 is not only sensitive to social predictions triggered by the videos but also to the specific emotional content.

### Prediction error effects on the late component N300 depend on emotional whole-body interaction

Next, we found an effect of prediction error on the processing of observed social interactions, as reflected by the N300, in line with our expectations and previous results relating the N300 to higher-order visual prediction errors (Chen et al., 2022). More specifically, enhancements of the N300 have been related to unexpected and violating conditions compared to expected and confirming conditions (Baker et al., 2023; Kumar et al., 2021; Truman & Mudrik, 2018). In line with these studies, we found a prediction error response (incongruent > congruent) for social interactions. Interestingly, this effect was only significant when the emotional reaction was neutral, indicating that neutral reactions may violate emotion predictions more strongly than angry ones.

These results indicate that the appropriateness of the reaction to an emotional interaction was extracted in the time window of the N300 (or 250-350ms post-stimulus onset) in our study. Unexpectedly, we found no effect of prediction strength on prediction-error responses in the N170 or N300, suggesting that these error responses do not necessarily depend on the availability of social predictions.

## 5 Conclusion

In sum, our results show that observing a social interaction generates perceptual predictions about how the behavior of the agents and these predictions affect cortical processing in the time window of the N170. The strength of this prediction effect measured at the final image is a function of how informative the preceding video is. This signifies that combined emotional expressions of interacting agents can be rapidly detected in early processing stages and that social interaction predictions influence information processing at perceptual and neural levels. Later prediction errors are reflected in the N300 amplitude, and this prediction error processing is most pronounced when observing a neutral bodily reaction. This suggests that later prediction may involve deeper cognitive processing reckoning with the emotional context in social interactions.

## Acknowledgements

This work was supported by the ERC Synergy grant (Grant agreement 856495; Relevance), by the Horizon 2020 Programme H2020-FETPROACT-2020-2 (Grant agreement 101017884 GuestXR) by the Research and Innovation Program H2020-EU.1.3.1 (Grant agreement 721385; Socrates), by the Horizon-CL4-2021-Human-01-21 (Grant agreement: 101070278; Re-Silence), and by China Scholarship Council (CSC202008440538).

## Supplementary

### Main effect of prediction error on N170 and N400

Analysis for main effects revealed a significant effect of *prediction error* (i.e., pooled across *reaction emotion* and *prediction strength*) on N170 and N400. More specifically, the incongruent condition elicited smaller N170 amplitudes and larger N400 amplitudes than the congruent condition (N170: incongruent: –0.51 ± 1.76 µV, congruent: –0.78 ± 1.90 µV, *F* (1, 27) = 7.13, *p* = 0.013, *η_p_^2^* = 0.21; N400: incongruent: –0.88 ± 1.36 µV, congruent: –0.71 ± 1.35 µV, *F* (1, 27) = 5.79, *p* = 0.023, *η_p_^2^* = 0.18).

**Fig. 5.**
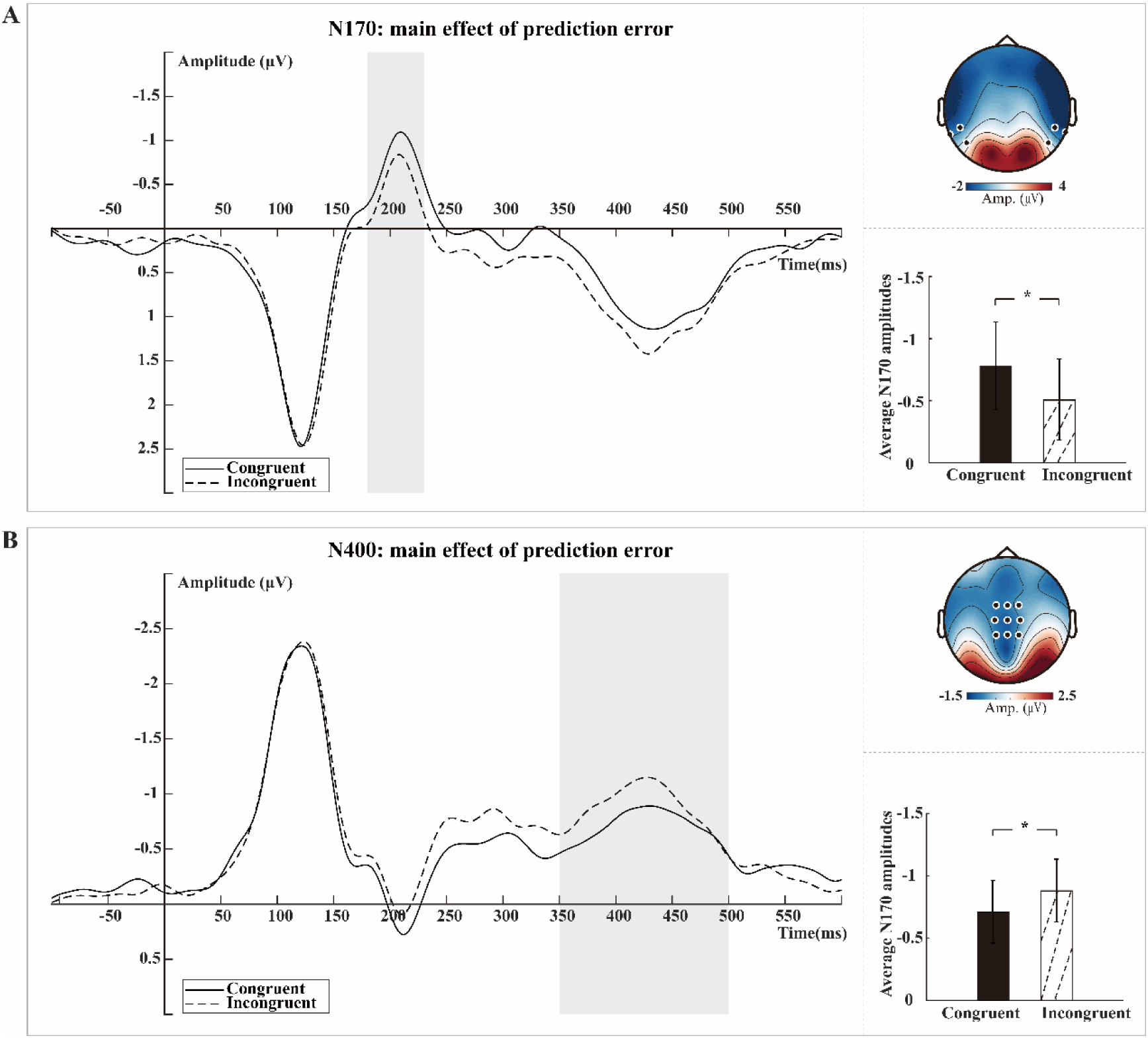
A-B. Main effect of *prediction error* on N170 and N400. Grand averaged ERPs are depicted per condition (congruent and incongruent) for N170 and N400 components separately (left). The shaded rectangle visualizes the time window (180-230ms for N170, and 350-450ms for N400) from which the average ERP amplitude was extracted. The highlighted black dots on the topographic map (right top) represent the electrodes from which the grand-averaged ERP for each component was extracted across all conditions. Bar plots (right bottom) illustrate the mean and SE across participants of each component’s amplitude per condition. *: *p* <0.05

### Main effect of prediction strength on N300 and N400

We further observed a significant main effect of *prediction strength* (i.e., pooled across *reaction emotion* and *prediction error*) on N300 and N400. More specifically, high prediction resulted in subsequent still images eliciting smaller N300 amplitudes and N400 amplitudes, compared with mid and especially low prediction (N300: high: –0.26 ± 1.51 µV, mid: –0.62 ± 1.74 µV, low: – 1.05 ± 1.33 µV, *F* (1, 27) = 9.54, *p* = 0.001, *η_p_^2^* = 0.26; N400: high: –0.26 ± 1.51 µV, mid: –0.84 ± 1.53 µV, low: –1.12 ± 1.31 µV, *F* (1, 27) = 10.13, *p* = 0.001, *η_p_^2^* = 0.27).

**Fig. 6.**
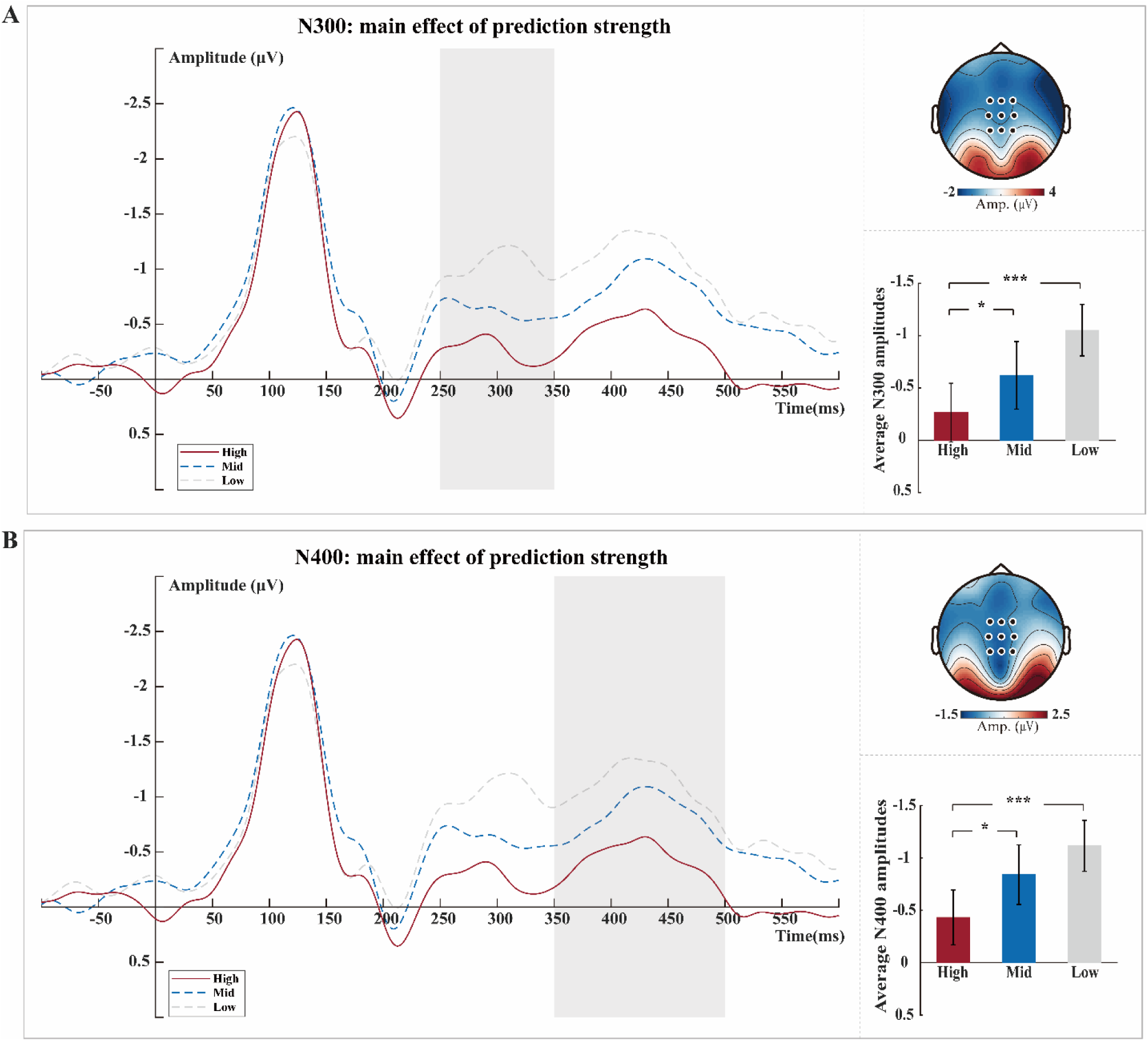
A-B. Main effect of *prediction strength* on N300 and N400. Grand averaged ERPs are depicted per condition (high, mid and low) for N300 and N400 components separately (left). The shaded rectangle visualizes the time window (250-350ms for N300, and 350-450ms for N400) from which the average ERP amplitude was extracted. The highlighted black dots on the topographic map (right top) represent the electrodes from which the grand-averaged ERP for each component was extracted across all conditions. Bar plots (right bottom) illustrate the mean and SE across participants of each component’s amplitude per condition. ***: *p* <0.001, *: *p* <0.05

